# Unidirectional migration of populations with Allee effect

**DOI:** 10.1101/2021.06.24.449708

**Authors:** Gergely Röst, AmirHosein Sadeghimanesh

**Affiliations:** Bolyai Institute, University of Szeged, Aradi vértanúk tere 1, Szeged, H-6720, Hungary

**Keywords:** population dynamics, migration, bifurcation, steady states, Allee effect, cylindrical algebraic decomposition

## Abstract

In this note we consider two populations living on identical patches, connected by unidirectional migration, and subject to strong Allee effect. We show that by increasing the migration rate, there are more bifurcation sequences than previous works showed. In particular, the number of steady states can change from 9 (small migration) to 3 (large migration) at a single bifurcation point, or via a sequences of bifurcations with the system having 9,7,5,3 or 9,7,9,3 steady states, depending on the Allee threshold. This is in contrast with the case of bidirectional migration, where the number of steady states always goes through the same bifurcation sequence of 9,5,3 steady states as we increase the migration rate, regardless of the value of the Allee threshold.

## 1 Introduction

Allee effect is a key concept in population ecology [1], and refers to the situation where a population has smaller growth rate at lower densities. In the case of strong Allee effect, there is a critical value called the Allee threshold, such that the population is declining whenever its density is below this threshold. The interplay of spatial dispersal and Allee effect is of special interest, and was subject of recent works [3–5, 8].

Figure 1 (a) in [5] illustrates how the structure of steady states change in a continuous time model of two connected populations living on identical patches with Allee effect. Generally, steady states collide in saddle-node bifurcations and disappear as we increase the migration rate between the patches, and the situation simplifies. However, it is not the case in a discrete time version, where attractors may appear and disappear in the presence of dispersal, as pointed out in [8]. In this note we show that steady states can appear even in the continuous time model as the migration rate increases, if we allow only one-way (unidirectional) migration between the patches. Here we completely describe the possible bifurcation sequences in the unidirectional case, which is more diverse than the bidirectional case. In particular, our results reveal that Figure 2 (a) in [5] is incomplete, and there are other possible routes via bifurcation sequences from 9 steady states (small migration) to 3 steady states (large migration). Our main tools are algebraic, and the computational files can be accessed here [6].

**Figure 1:**
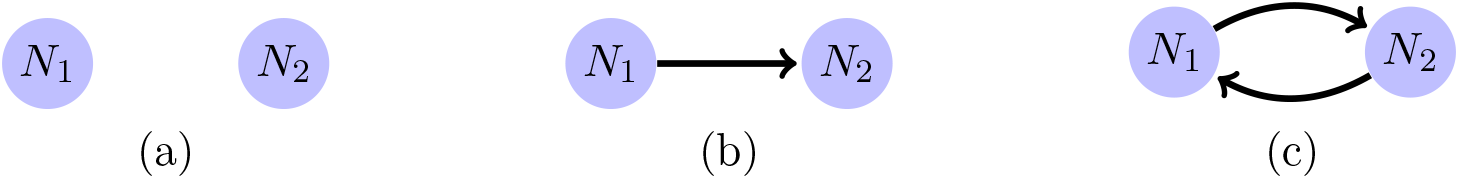
There are only three non-isomorphic digraphs on two nodes. (a) Two isolated nodes. (b) There is one directed edge from one of the two nodes to the other one. (c) There are two directed edges between the two nodes.

**Figure 2:**
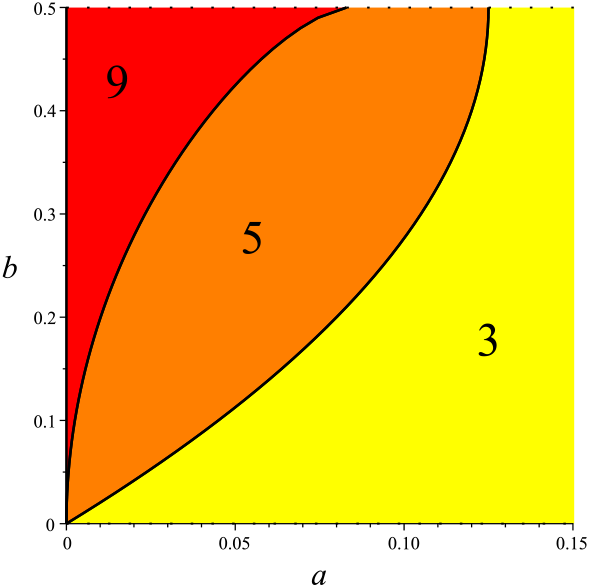
The parameter region 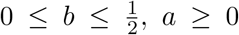 of the 2 patches model (1) with the connectivity graph as shown in Figure 1c is partitioned to sub-regions with invariant number of steady states. The number of steady states is written on each open region.

## 2 Two patch model

We consider a two patches population model with strong Allee effect and spatial dispersal and different possible connectivity of the two patches. There exist three non-isomorphic digraphs with two nodes; two disconnected nodes, a directed graph with a source and a target, and finally a fully connected digraph. These three digraphs are shown in Figure 1. Let *N_i_* denote the population at patch no. *i* with carrying capacity 1, let *b* ∈ (0, 1) be the Allee threshold, and *a* ≥ 0 be the spatial dispersal rate. Then the population of the patches is modeled by the following system

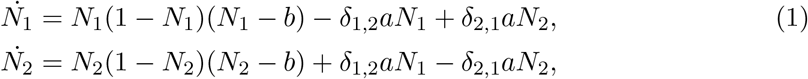

where *δ_i,j_* is 1 if there is a directed edge from patch *i* to patch *j*, and 0 otherwise. This is the same as [5, Equation *M_α_*] for *r* = 2 and *g_i_*(*N_i_*) = (1 − *N_i_*)(*N_i_* − *b*), and when the connectivity graph is fully connected is the same as [7, Equation 2.1] for *n* = 2.

For a fixed choice of parameter values, a steady state or equilibrium of system (1) is a non-negative solution to the system of equations made by letting 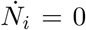 for *i* = 1, 2. Therefore to study the number of steady states of our model, the following parametric system of polynomial equations needs to be studied:

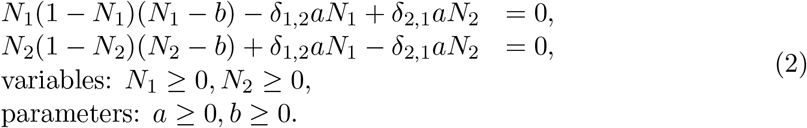

We investigated this parametric system using `RootFinding[Parametric]` library in Maple [2]. In contrast with the case *n* = 3 at [7], the case *n* = 2 is not heavy for a simple computer to handle (computations performed on Windows 10, Intel(R) Core(TM) i7-2670QM CPU @ 2.20GHz 2.20 GHz, x64-based processor, 6.00GB (RAM)). Thus without using any numerical algorithm one can get an exact description of the boundaries of different parameter regions where the system (2) has different number of non-negative solutions.

It is clear that in the absence of the spatial dispersal between the two patches, the system has 9 non-negative steady states for *b* ∈ (0, 1). In the sequel we concentrate on the two other connectivity cases shown in Figure 1b and 1c.

## 3 Bidirectional (both ways) migration

Consider first the connectivity graph in Figure 1c. In this case *δ*_1,2_ = *δ*_2,1_ = 1. By Lemmas 2.1 and 2.2 from [7], it is enough to restrict the parameter space to 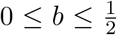. Figure 2 shows the decomposition of the the parameter region 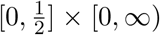 with respect to the number of steady states. Fix a value for *b*, say *b*\*. A value *a* = *a*\* is said critical for *a* if the point (*a*\*, *b*\*) is on one of the boundaries between different regions. For any *b* ≠ 0, there exists two critical values for *a*. The sequences of the number of steady states for a fixed value of *b* ≠ 0 when increasing *a* (considering only open regions) are always the same as listed below:

• 9, 5, 3.

When *a* = 0, there are 9 steady states, passing the first critical value of *a*, two pairs of steady states collide and disappear in saddle node bifurcation events. Passing the second critical value, two other steady states meet at the middle steady state and the two disappear in a pitchfork bifurcation event. This has been shown in [5, Figure 1a] where *b* = 0.3. Indeed in this case, the two connected populations has a simple behavior as was initially guessed.

## 4 Unidirectional (one-way) migration

Consider next the connectivity graph in Figure 1b. In this case *δ*_1,2_ = 1, but *δ*_2,1_ = 0. Here only [7, Lemma 2.1] holds and not [7, Lemma 2.2]. Thus this time the parameter region necessary to study is 0 ≤ *b* ≤ 1. In contrast with the previous case, the number of critical values for *a* are not always the same for fixed values of *b*. We refer to the values of *b* where the number of critical values for *a* changes as critical values for *b*. We found that the critical values for *b* are 0, 1, and *β*_1_, and *β*_2_, where *β*_2_ = 1/2, and *β*_1_ is an algebraic number, specifically the smallest positive real root of 59*b*^4^ − 16*b*^3^ − 214*b*^2^ − 16*b* + 59. With seven digits accuracy *β*_1_ ≃ 0.4961661.

The sequences of the number of steady states for a fixed value of *b* when increasing *a* (considering only open regions) are listed below;

- 9, 7, 5, 3.
- 9, 7, 9, 3.
- 9, 3.

Figure 3a shows the decomposition of the parameter space with respect to the number of steady states. The region where the increment in the number of steady states happens is enlarged at Figure 3b. See also Table 1 for the detailed description of the parameter regions with different number of steady states. Figure 2a in [5] has shown the sequence of steady states for *b* = 0.3 where the temporary increment in the number of steady states does not happen.

**Figure 3:**
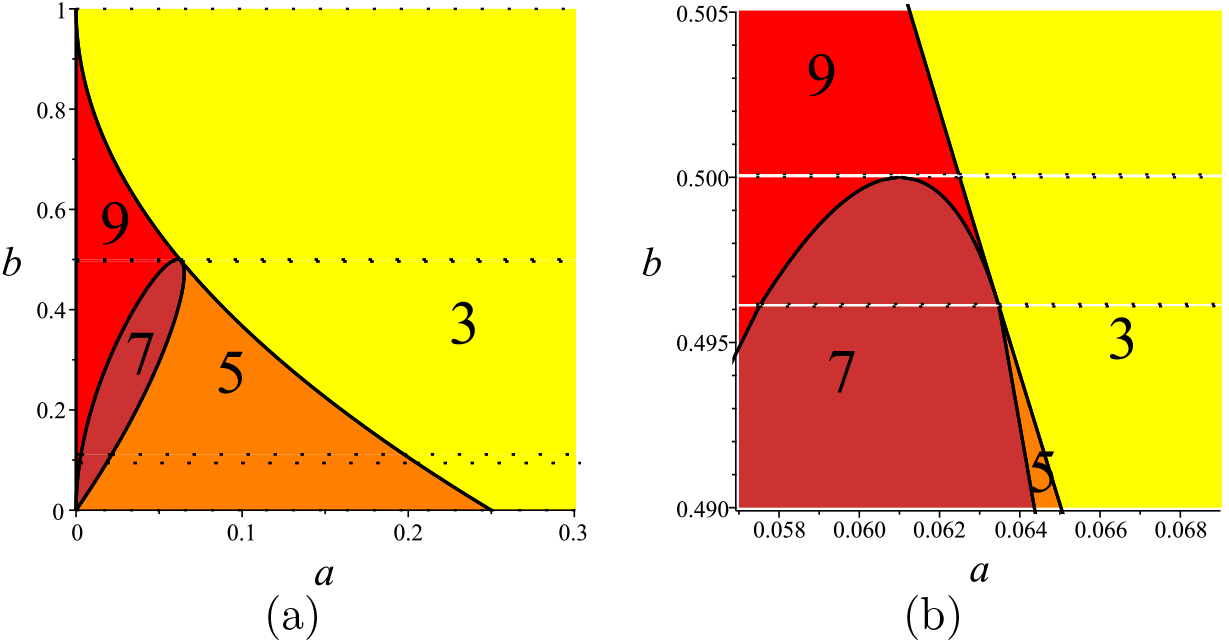
(a) The parameter region 0 ≤ *b* ≤ 1, *a* ≥ 0 of the 2 patches model (1) with the connectivity graph as shown in Firgure 1b is partitioned to sub-regions with invariant number of steady states. (b) When *β*_1_ < *b* < 0.5 the number of steady state temporary increases when increasing *a*, this region is the tiny red colored part between the brown and yellow regions.

**Table 1:**
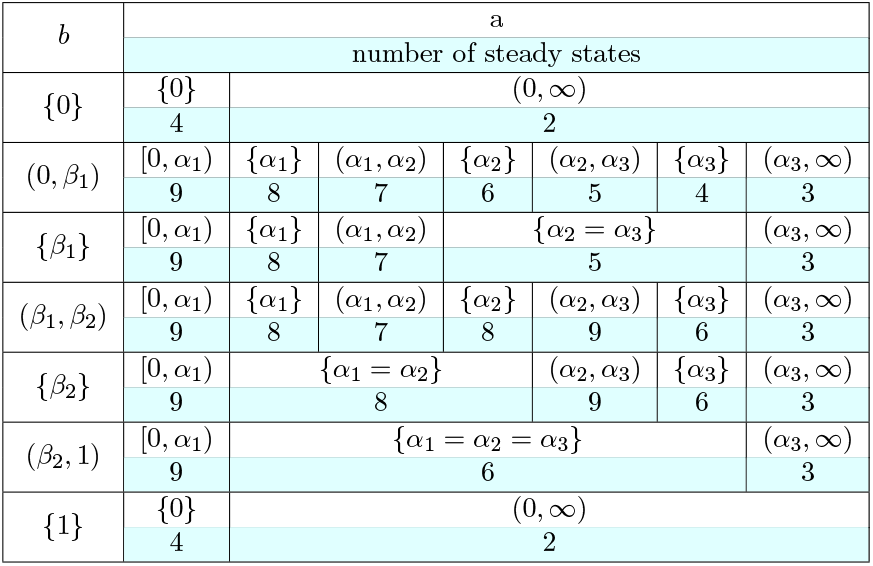
The number of non-negative steady states of the the 2 patches model (1) with the connectivity graph as shown in Firgure 1b for the parameters *a* ≥ 0, 0 ≤ *b* ≤ 1. The first column shows the interval of *b*. Each row for the intervals of *b* has two sub-rows, the first sub-row shows the interval of *a* and the second sub-row states the number of steady states.

Figure 4 shows the five possible sequences for *b* ≠ 0, 1 (including the cases *b* = *β*_1_ and *b* = 0.5) together with the stability of the steady states. Figure 5 is a schematic figure simplifying the behavior of the system for these 5 cases. All bifurcation events here are saddle-node except at two points. At *b* = 0.5, by increasing *a* to cross its first critical value, a transcritical bifurcation event happens where two steady states meet and then continue their paths. At *b* = *β*_1_, by increasing *a* to cross its second critical value, two steady states that already had met and left the real plane return at exactly the location where another pair of steady states are meeting to leave the real plane. Therefore at this moment, there are four steady states at a single point.

**Figure 4:**
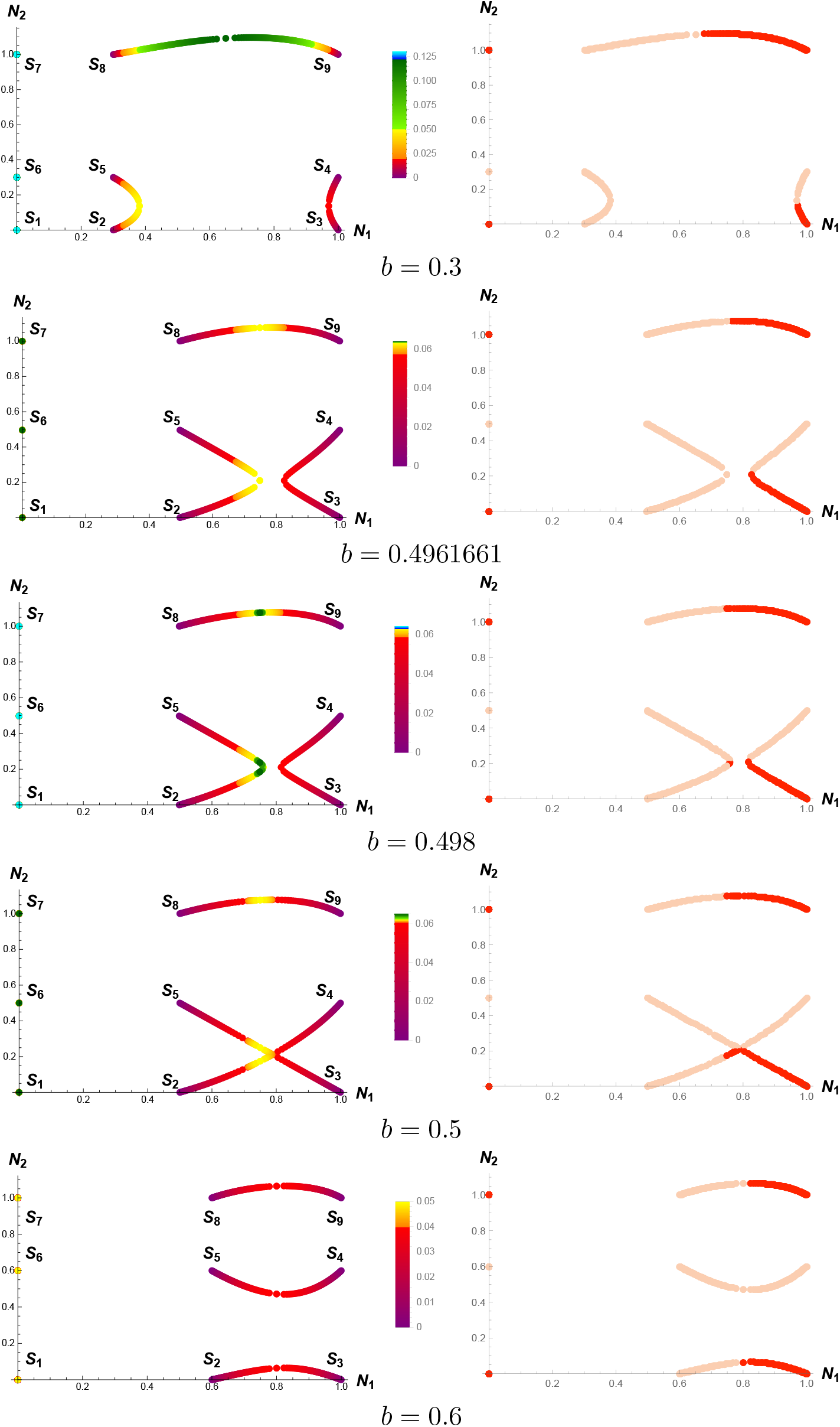
Sequences of the steady states for fixed values of *b*, but varying values of *a*. In the left side the steady state points are colored with respect to the value of *a* that can be read from the color bar next to the plots. In the right side the steady states are colored with respect to their stability. Stable steady states are colored by red, whereas the unstable ones are colored by pink.

**Figure 5:**
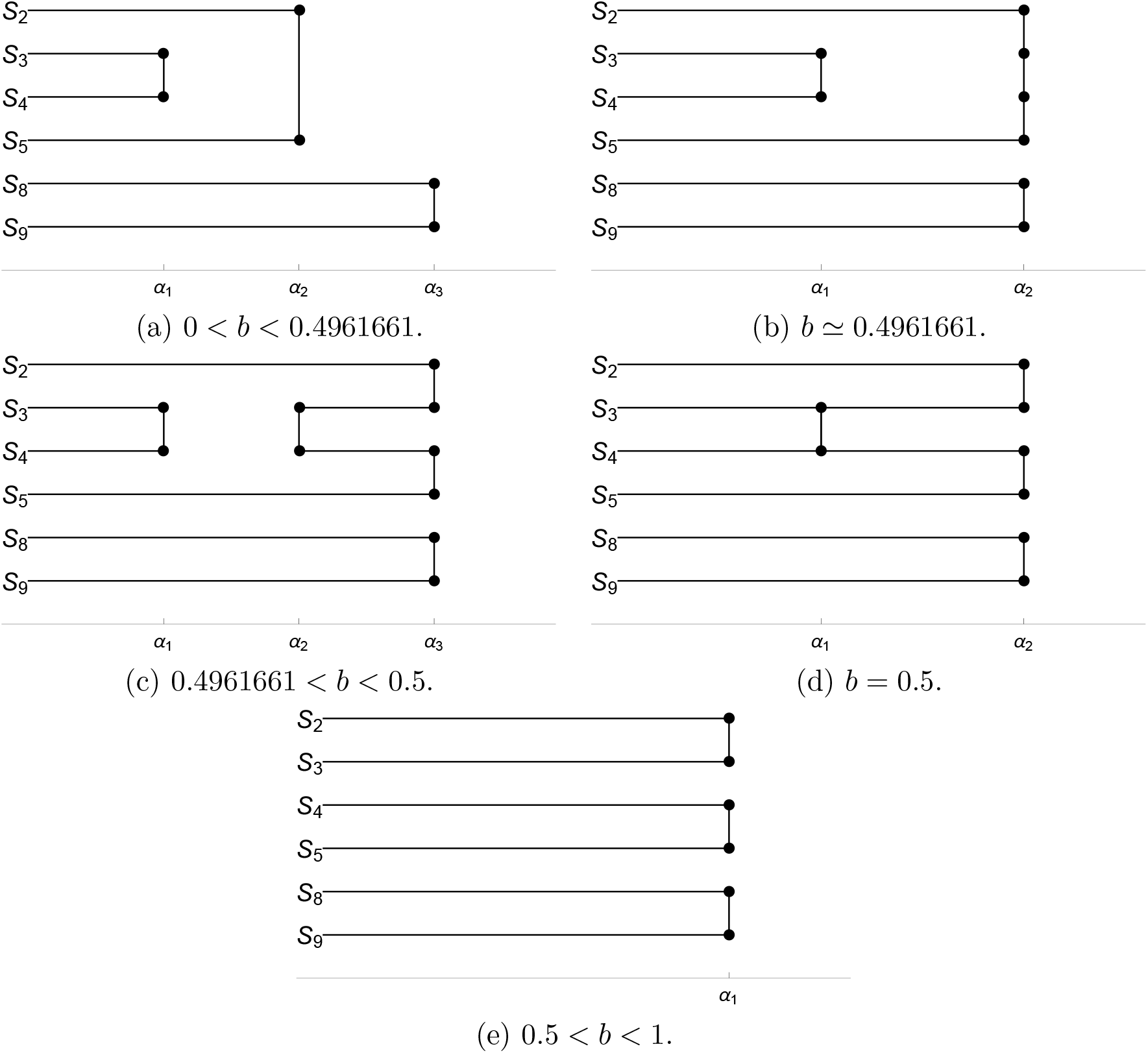
Schematic plots showing when and which of the initial 9 steady states of the 2 patches model (1) with the connectivity graph Figure 1b collide and disappear. The initial nine steady states for *a* = 0 are shown at Figure 4 which are *S*_1_ = (0, 0), *S*_2_ = (*b,* 0), *S*_3_ = (1, 0), *S*_4_ = (1*, b*), *S*_5_ = (*b, b*), *S*_6_ = (0*, b*), *S*_7_ = (0, 1), *S*_8_ = (*b,* 1) and *S*_6_ = (1, 1). In contrast with the initial guess that by increasing *a*, the steady states only meet and disappear, as one can see in (c) two steady states that previously have met and went out of the real plane, can meet again and return from the non-real complex plane back to the real plane. The horizontal line shows the values of *a* which is 0 at the left and increases as we move to the right. All bifurcation events are of the saddle-node type, except at *a* = *α*_2_ in part (b) and at *a* = *α*_1_ in part (d). In the first case, the two steady states *S*_3_ and *S*_4_ are coming back at the same location where *S*_2_ and *S*_5_ are meeting and then all four of them leave the real plane. At the second case, a transcritical bifurcation occurs. *S*_3_ and *S*_4_ meet and then continue their paths.

## 5 Discussion

In [5] it was stated that for small enough dispersal rate, the *n* patches model has 3^*n*^ steady states and generally the situation simplifies when migration is larger, eventually leading to only 3 steady states resembling a simple one larger patch consisting of all populations together. It was expected to see a monotone decrease in the number of steady states by increasing the dispersal rate until in [8] for the discrete version of this model, and in [7] for *n* = 3 it was shown that the behaviour can be more complex than it might look at an initial thought. The authors of this paper has completely classified all possible bifurcations for the case *n* = 3 in [7] and have shown that a temporary increment in the number of steady states can occur for specific choices of the Allee threshold, provided that all patches are connected both ways. Here we proved that we can see an increment in the number of steady states even for the case *n* = 2, if we allow only one-way migration. We confirm that for bidirectional migration, there is only one bifurcation sequence resulting in 9,5,3 steady states as the migration rate increases. However, if the migration is only one-way, there is a variety of bifurcation sequences that we fully classify in Table 1, Figure 4, and Figure 5.

## Acknowledgements

Gergely Röst was supported by Hungarian grants EFOP-3.6.1-16-2016-00008, NKFIH FK 124016, and TUDFO/47138-1/2019-ITM7. AmirHosein Sadeghimanesh was funded by NKFIH KKP 129877.

